# RESCUE: recovery of unattributed expression patterns in spatial transcriptomics

**DOI:** 10.1101/2025.08.11.669542

**Authors:** Young Joo Lee, Seokjin Yeo, Alex W. Schrader, Ian M. Traniello, Amy Cash Ahmed, Gene E. Robinson, Hee-Sun Han, Sihai Dave Zhao

## Abstract

Spatial transcriptomics (ST) enables gene expression profiling while preserving the spatial architecture of intact tissue. Analyzing ST data often proceeds by first extracting cell-level information, typically through cell segmentation or cell-type deconvolution. However, a critical oversight has been that a substantial portion of molecular expression is systematically lost or unannotated by these methods. This lost expression can arise from diverse and biologically important sources like fragile or underrepresented cell types, subcellular structures like neurites, and extracellular expression. These omissions can result in biased analyses and incorrect or incomplete biological interpretations. We describe a new computational method, RESCUE, that can recover the unattributed spatial expression patterns missed by existing ST analysis methods and enable robust inference even when reference is incomplete. We validate RESCUE using MERFISH data from the honey bee brain and apply it to multiple ST datasets to demonstrate how it can reveal novel insights into complex tissue biology.

## INTRODUCTION

Profiling cellular expression within its spatial context has become possible with the recent development of spatial transcriptomics (ST) technologies, which measures gene expression in intact tissue sections^1,2^. Analyzing ST data typically proceeds by extracting cell-level information from the spatial expression data, as cells are natural units of analysis. With high-resolution platforms such as MERFISH^3^, 10x Visium HD^4^, or 10x Xenium^5^, this is usually done using cell segmentation^6–8^. With low-resolution platforms like 10x Visium^9^ and Slide-seq^10,11^, where each spatial location can contain multiple cells, it is common to deconvolve the observed expression into separate contributions from different cell types, based on reference single-cell/nuclei RNA-seq (sc/snRNA-seq) data^12–16^.

However, a major problem with these existing ST analysis methods is that they can fail to capture important expression patterns. First, they may omit entire cells or cell types. Cell segmentation algorithms for high-resolution data have difficulty with cells with complex morphologies, such as fibroblasts, or cells lacking nuclei, like erythrocytes^17,18^. Deconvolution methods for low-resolution data cannot account for cell types that are absent from the reference sc/snRNA-seq data because they are too fragile to survive tissue dissociation^19–21^. Second, existing ST approaches can overlook expression patterns unique to cellular substructures. Segmentation methods simply drop transcripts located in substructures that are difficult to segment, such as the axons and dendrites of neurons. These fragile structures are also typically destroyed during tissue dissociation and therefore are not represented in reference single-cell data used in deconvolution. Indeed, it has been estimated that over 40% of the total transcriptome of an adult mouse brain may be missed by sc/snRNA-seq^22^. Similarly, reference snRNA-seq data, by definition, ignores cytoplasmic expression. Finally, existing methods do not capture extracellular tissue expression patterns, which cannot be studied by segmentation or deconvolution.

These difficult-to-capture expression patterns can be critically important. First, they can bias analyses that do not take them into account. An example with dramatic implications is that deconvolution estimates of cell-type proportions can be invalid if the reference is missing any cell types^23^, as we illustrate later in this work. Second, there may be important biological processes that are unique to the overlooked expression patterns. For example, important missed cell types include adipocytes^5^, which play crucial roles in metabolic disease and cancer^24^, and neutrophils^25^, which are core components of the immune system. Important missed substructural expression includes axonal or dendritic transcripts that are central to synaptic development and plasticity^26–30^ and diseases such as amyotrophic lateral sclerosis^31,32^, as well as cytoplasmic transcripts that regulate key processes such as metabolism, signaling, and stress responses^33–35^ and are involved in Alzheimer’s disease^36,37^. Important missed extracellular expression patterns include extracellular RNAs that mediate intercellular communication in cancer^38–40^.

We describe a new computational method, RESCUE, that can recover the spatial expression patterns missed by single-cell inference on ST data. Our approach is based on making a fundamental correction to the class of statistical models currently used for integrating ST data and reference data, such as sc/snRNA or cell segmentation results. Briefly, our method partitions observed gene expression into one component explainable by the reference data, which we refer to as “canonical” expression, and another component that is not, which we refer to as “unattributed” expression. Our purified canonical ST expression component can be used to correct existing cell-type deconvolution methods. Our recovered unattributed ST expression component gives us the new ability to specifically target biology that is distinct from our reference data, as discussed above. The unattributed expression recovery step in RESCUE is analogous to negative selection in experimental biology^41–44^, where we identify and remove canonical expression. We demonstrate that RESCUE accurately recovers unattributed expression, enables robust deconvolution of ST data despite incomplete reference data, and reveals biologically meaningful patterns overlooked by existing methods. While we focus here on ST data, we note that RESCUE can also be directly applied to non-spatial modalities, such as bulk RNA-seq data.

## RESULTS

### Overview of RESCUE

RESCUE is predicated on a new model that we propose for integrating ST and reference single-cell data, and can be applied to diverse ST platforms, including MERFISH and Visium, across different organisms. Most existing models^12–16^ posit that ST expression at each location is composed of a linear combination of expression patterns present in the reference, where the coefficients of this linear combination are interpreted as the proportions of each reference cell type in that location. However, as described in the introduction, there are numerous sources of ST expression that are left out of this model; we will refer to these as unattributed expression patterns, because they are not attributed to canonical expression patterns reflected in the reference.

To address this issue, we propose a corrected model (**Fig. 1**) that allows the ST profile to have both canonical and unattributed expression components. While it appears minor, this change has important implications. Under this model, results from the statistics literatures^45–47^ imply that under certain conditions it is possible to simultaneously recover unattributed expression and estimate the linear combination of reference patterns that constitute canonical expression. The conditions are that only relatively few genes show non-zero, or “sparse” expression, in the unattributed component, which is reasonable given the nature of unattributed expression and the read depth/detection efficiency of ST technologies (**Methods**).

**Figure 1.**
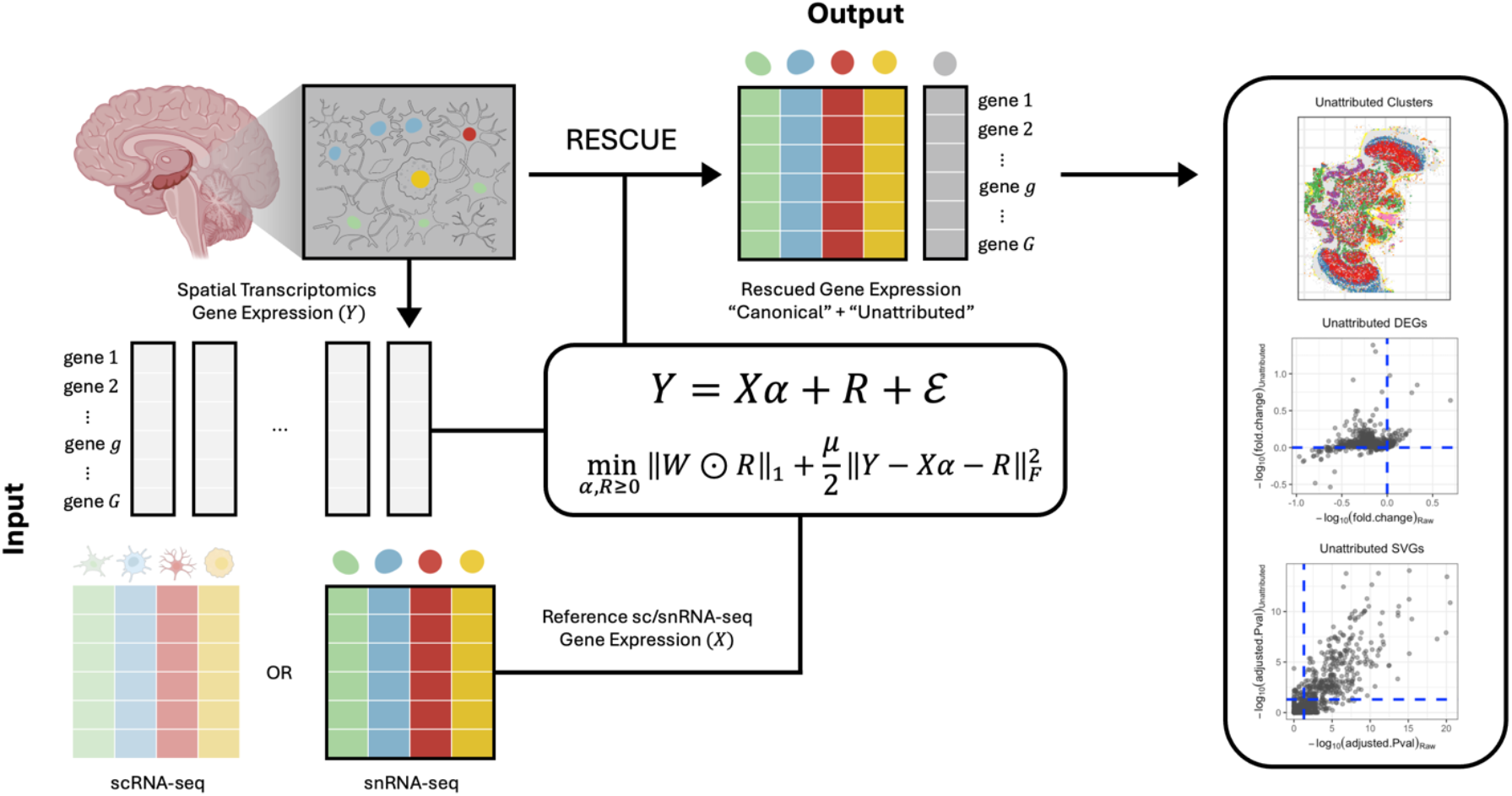
Schematic overview of RESCUE. RESCUE is a novel computational negative selection method designed to recover the spatial expression patterns missed by standard single-cell inference on ST data. It takes as input ST data to be decomposed, along with a reference matrix constructed from sc/snRNA-seq data from the same tissue, and performs robust sparse recovery via a regularized multivariate regression framework. RESCUE partitions observed gene expression into a reference-explainable “canonical” component and a residual “unattributed” component. As a result, the purified canonical expression enables more accurate deconvolution, while the recovered unattributed expression enables downstream analyses such as clustering, differentially expressed gene (DEG), and spatially variable gene (SVG) analyses. Created with BioRender.com.

Leveraging this insight, we developed RESCUE, an algorithm that simultaneously deconvolves ST data while rescuing spatial unattributed expression. RESCUE solves a regularized robust multivariate regression problem and can be applied to both low- and high-resolution ST data. Unlike existing deconvolution methods^12–16^, it is robust to incomplete references, such as snRNA-seq data which lack cytoplasmic expression profiles. We emphasize that the interpretation of the unattributed component depends on both the biological context of the tissue and the choice of the reference dataset (**Methods**). As a result, the unattributed component may comprise multiple biological sources, including expression from cell types missing in the reference, substructural expression, and extracellular RNA. RESCUE is designed to recover these signals jointly rather than to assign them to predefined categories. The rescued unattributed expression can then be further decomposed using standard unsupervised approaches, such as clustering and non-negative matrix factorization (NMF), to identify distinct expression patterns within the unattributed component. In addition, differentially expressed gene (DEG) and spatially variable gene (SVG) analyses can be applied to characterize the biological and spatial features of the recovered unattributed signals. Careful consideration of biological structures and tissue organization is therefore essential when interpreting the unattributed expression. We illustrate this in a broad range of examples below.

### Simulations

To evaluate the performance of RESCUE, we conducted simulation studies on synthetic data designed to mimic real ST measurements with varying degrees of unattributed expression. We simulated ST data using real scRNA-seq data from the honey bee brain^48^ by introducing synthetic unattributed expression of a small number of genes (**Methods**). Unlike other tissues where cell bodies and non-cell-body structures are spatially intermingled, honey bee brains provide a unique benefit for simulating unattributed expression due to the existence of the neuropil region, where axons and dendrites are spatially segregated from cell bodies. Molecular expression in these regions is largely absent in single-cell sequencing data due to the structural fragility of neurites. During the tissue dissociation step, a necessary step in single-cell sequencing, neurites are mostly sheared off and lost. In contrast, ST measurements performed on intact tissue sections preserve these structures and therefore capture molecular expression within neuropils.

To guide the choice of sparsity levels used in our simulations, we examined empirical expression patterns in the neuropil region of our in-house MERFISH data. We found that neuropil regions exhibit highly sparse gene expression, with an average sparsity level of 0.92 across spatial locations (**Fig. 2a**). Here, sparsity refers to the structure of the unattributed component (i.e., that only a subset of genes is present at a given spatial location), rather than to low expression levels (**Methods**). Based on this empirical distribution, we simulated unattributed expression across a range of sparsity levels by varying the proportion of zero entries (ranging from 0.8 to 0.95). In addition, we varied the proportion of unattributed expression (ranging from 0 to 0.8) to cover different levels of the overall magnitude of unattributed expression. Importantly, these parameter ranges fall within those observed in real biological samples and are not artificially chosen to favor RESCUE.

**Figure 2.**
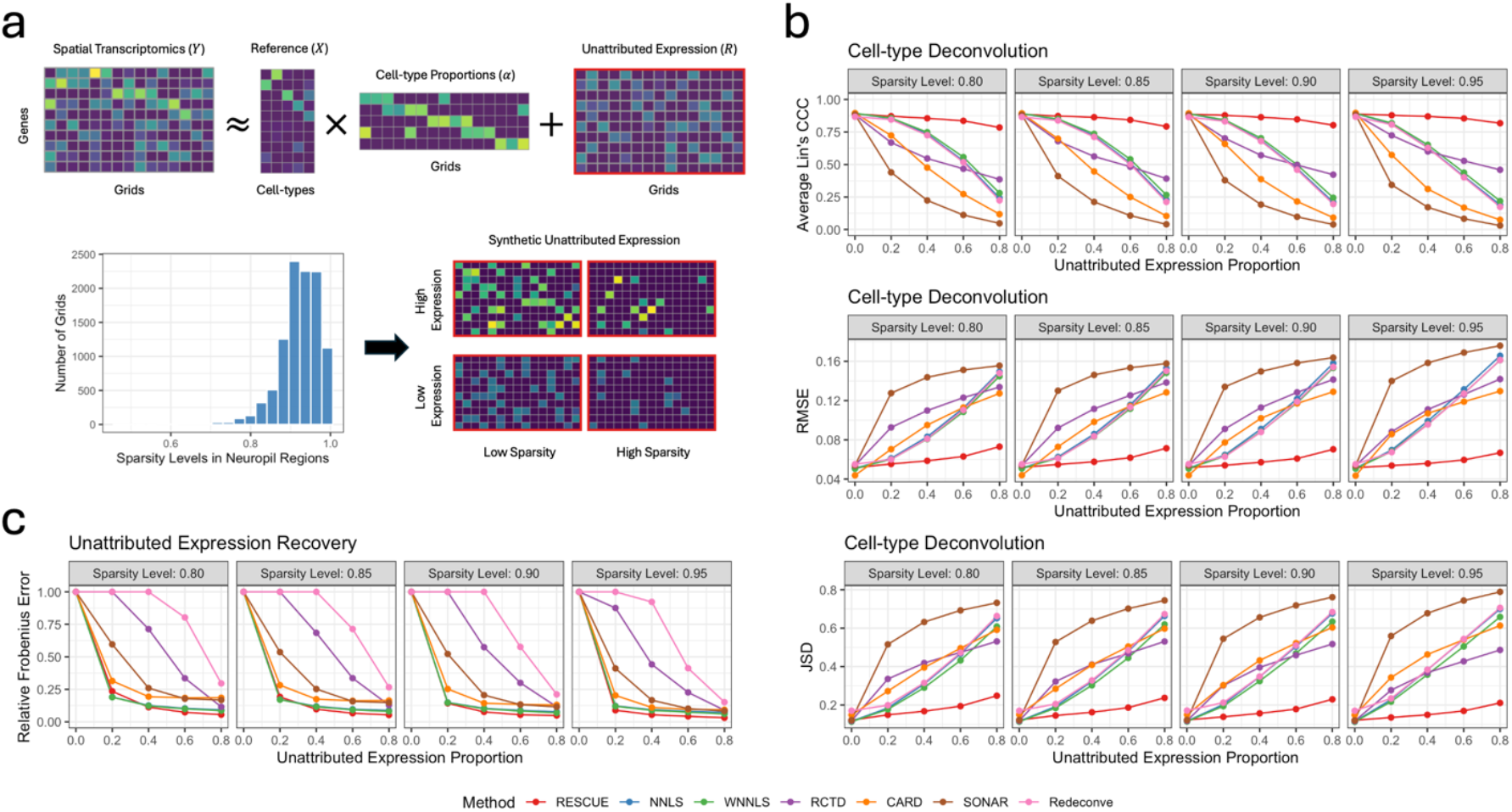
Simulation results. **a**. Schematic of the simulation framework. ST data are generated as the sum of reference-driven canonical expression and synthetic unattributed expression. The distribution of sparsity levels observed in neuropil regions of the honey bee brain motivates the simulated sparsity regimes. Representative examples of synthetic unattributed expression under low- and high-sparsity/expression levels are shown. **b**. Cell-type deconvolution performance across methods, evaluated using average Lin’s concordance correlation coefficient (CCC; top), root mean squared error (RMSE; middle), and Jensen-Shannon divergence (JSD; bottom) as a function of the sparsity/proportion levels of the unattributed expression. **c**. Unattributed expression recovery performance measured by relative Frobenius error under the same simulation settings.

We first compared the deconvolution performance of RESCUE with existing methods. **Fig. 2b** summarizes the accuracy of cell-type proportion estimates across methods, measured using the average Lin’s concordance correlation coefficient^49^ (CCC), root mean squared error (RMSE), and Jensen-Shannon divergence (JSD) (**Methods**). As expected, the performance of standard deconvolution methods deteriorates sharply as the level of unattributed expression increases. In contrast, RESCUE maintains high accuracy across all conditions, demonstrating robustness to both high unattributed content and reduced sparsity. These results indicate that failing to explicitly model unattributed expression can lead to systematic biases in cell-type proportion estimates.

We also evaluated the accuracy of unattributed expression recovery (**Fig. 2c**) using the relative Frobenius error between the true and estimated unattributed expression matrices (**Methods**). Overall, RESCUE achieves the lowest recovery error even under high unattributed expression and low sparsity levels, whereas other methods deviate substantially from the ground truth, especially when unattributed signals become strong. These results indicate that explicitly modeling unattributed expression enables accurate recovery of hidden transcriptional signals across a wide range of sparsity and signal-strength regimes.

To assess robustness and generalizability, we expanded the simulation study by varying noise levels via library size variability across spots and varying the effective grid resolution via changes in the average total UMI count per spot (**Supplementary Fig. 1**). As expected, overall performance tends to decrease with increasing library size variability and increase with higher average total UMIs. Across these regimes, RESCUE remains among the top-performing methods, demonstrating robustness across a range of noise levels and grid sizes.

### RESCUE accurately recovers the unattributed transcriptome in real tissues

Using a custom MERFISH platform^50,51^, we generated single molecule-resolution ST data from coronal sections of a honey bee brain, targeting 130 genes relevant to aggression (**Supplementary Table S1**)^52^. Transcripts in axons and dendrites, which are collectively termed as neurites, have been difficult to profile due to segmentation challenges. Cell segmentation methods, such as Cellpose^6^ and Baysor^7^, can only identify the boundaries of neuronal cell bodies, also called somata. Although individual axons and dendrites can be stained in 2D cell cultures^53^, in native tissues they extend across multiple imaging planes and therefore cannot be reliably segmented as individual cellular units. As a result, neurite-associated expression has traditionally been ignored or incorrectly merged into nearby cytosolic expression.

We first experimentally validated that RESCUE can accurately recover gene expression in the neuropil region of the honey bee brain (**Fig. 3a**). This validation is possible because the physical separation of neurites and somata in the brain makes it simple for existing algorithms to separate somatic from unattributed expression. Cell soma were segmented using Cellpose^6^, revealing that only 73.5% of transcripts were localized within cells, with the remaining 26.5% comprising the unattributed expression. Using honey bee brain scRNA-seq data^48^ as a reference, we then applied RESCUE and competing methods to our MERFISH data after dividing the brain into grids of size 20 × 20 *μm*^2^ (**Methods**). For each grid, we estimated the proportion of canonical expression relative to total expression and obtained a ground-truth estimate by identifying transcripts localized within cell bodies via cell segmentation. RESCUE’s estimates closely matched the ground truth (**Fig. 3b**) and were more accurate than those from other methods, as quantified by Lin’s CCC and RMSE (**Methods**). Notably, RESCUE relies only on reference scRNA-seq data and does not use cell segmentation information.

**Figure 3.**
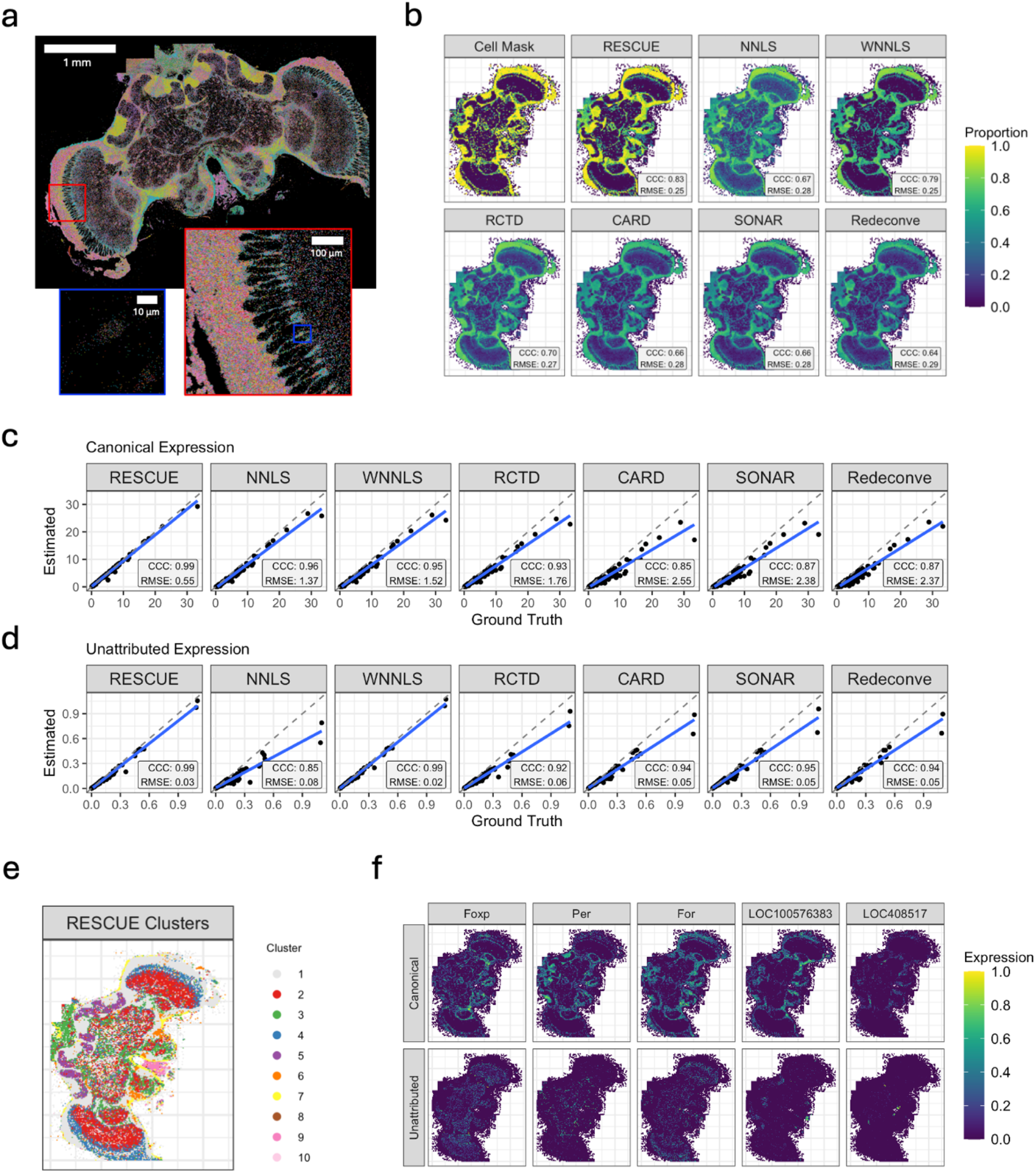
Application to the honey bee brain MERFISH data. **a**. MERFISH profiling of a coronal honey bee brain section. Each dot represents an individual transcript, with genes shown in distinct colors. A substantial fraction of transcripts localize to neurites, rather than to cell bodies. **b**. Comparison of segmentation-based ground-truth (“Cell Mask”) and estimated somatic proportions. Cell mask ratio is computed as the proportion of MERFISH spots in each grid lying within a cell. Lin’s concordance correlation coefficient (CCC) and root mean squared error (RMSE) between ground-truth and estimated proportions are reported. **c**. Comparison of estimated versus ground-truth canonical gene expression levels in grids corresponding to pure cell body regions. **d**. Comparison of estimated versus ground-truth unattributed gene expression levels in neuropil-only grids lacking segmented somata. For **c** and **d**, each point represents a gene averaged across grids; grey dashed lines indicate the identity line, and blue lines denote linear regression fits. Lin’s CCC and RMSE between ground-truth and estimated average expression are reported. **e**. Clustering of the unattributed expression profiles inferred by RESCUE reveals distinct transcriptional niches within neuropil regions, while somatic regions collapse into a single cluster. **f**. Spatial expression patterns of representative genes, shown separately for canonical and unattributed components. Expression values are min-max normalized separately for each panel.

To further clarify the source of discrepancies among methods, we evaluated canonical and unattributed expression separately in anatomically well-defined regions. Specifically, we examined grids corresponding to pure cell bodies and compared the average canonical expression of each gene against the ground truth derived from segmentation (**Fig. 3c**). RESCUE closely matched the true somatic expression levels, whereas competing methods systematically underestimated canonical expression. This behavior reflects the fact that existing deconvolution approaches are designed to estimate cell-type proportions rather than expression levels, and under the subtraction-based framework (**Methods**), expression that is not well explained by the reference profiles is left as residual rather than attributed to the canonical component. Conversely, in grids containing no identifiable cell bodies, RESCUE accurately recovered unattributed expression in neuropil-only regions (**Fig. 3d**), while competing methods were significantly less accurate. Together, these results demonstrate that RESCUE effectively partitions the full transcriptome into canonical and unattributed components in real tissue, capturing valuable expression signals that would otherwise be lost using conventional approaches.

Importantly, transcripts outside segmented somata reflect genuine biological signals rather than technical artifacts: in our MERFISH sample (**Fig. 3a**), neuropil-associated transcripts form structured spatial patterns that are clearly segregated from DAPI-stained nuclei (**Supplementary Fig. 2**). Moreover, the MERFISH protocol includes gel embedding, expansion, and clearing steps that reduce autofluorescence and suppress nonspecific signal^54,55^. These signals, which would have been ignored by standard cell segmentation approaches, highlight the importance of RESCUE in capturing overlooked transcriptomic information.

Finally, to investigate whether the unattributed expression recovered by RESCUE reflects structured biological organization rather than unstructured noise, we performed clustering analysis of the unattributed expression profiles across spatial locations. Clustering revealed anatomically distinct transcriptional niches within neuropil regions (**Fig. 3e**), indicating that the recovered unattributed component exhibits coherent spatial patterns. Specifically, distinct clusters localize to known neuropil structures, including the mushroom body calyces, antennal lobes, optic lobes, and regions proximal to the central complex. In contrast, spatial locations corresponding to somatic regions collapsed into a single cluster, consistent with the fact that canonical expression is explicitly removed, and only minimal unattributed signal remains in these regions. Examination of representative genes within the neuropil-associated clusters showed spatially structured expression patterns that are largely absent from the canonical component (**Fig. 3f**). These findings further support that the unattributed expression recovered by RESCUE captures biologically meaningful and spatially organized signals, revealing hidden tissue structure that is not accessible through conventional analyses.

### RESCUE enables accurate deconvolution of ST data with incomplete reference

Here, we illustrate how RESCUE’s estimated canonical expression component can be used to achieve accurate deconvolution of ST data when key cell types are missing from the reference, a challenge for all existing methods that has been noted in a recent study^23^. We analyzed paired single-cell FFPE-seq (scFFPE-seq) and 10x Visium ST data from human breast cancer tissue^5^. In this ST dataset, adipocytes are manually annotated in approximately 10% of spatial spots (**Fig. 4a**). However, adipocytes are absent from the scFFPE-seq reference, likely due to their high lipid content interfering with fixation and processing. These cells are prone to floating, rupturing, or sticking to plastic surfaces during tissue dissociation, which hinders their recovery in single-cell sequencing protocols^5,17^. As a result, reference-based methods that assume the observed expression is fully explained by the available reference cell types tend to misattribute adipocyte-derived transcripts to other cell types.

**Figure 4.**
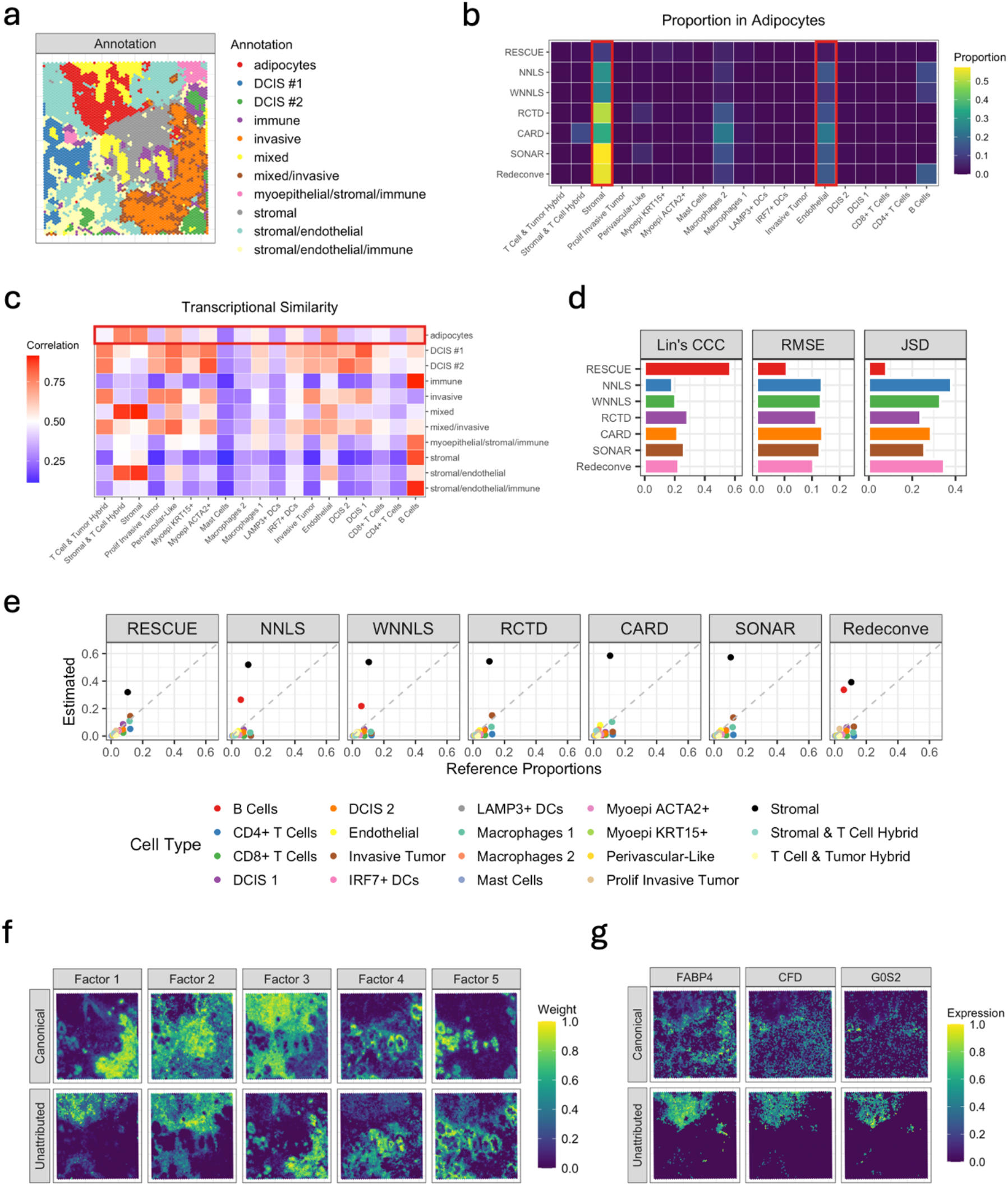
Application to the human breast cancer Visium data. **a**. Manual annotation of Visium ST data from human breast cancer tissue. **b**. Estimated cell-type proportions in adipocyte regions across methods. Red box highlights that existing deconvolution methods misattribute adipocyte-derived signal to other reference cell types due to the absence of adipocytes in the reference, whereas RESCUE mitigates such misassignment by modeling unattributed component. **c**. Heatmap of correlations between pseud-bulk Visium ST and scFFPE-seq reference cell types. Red box highlights high correlations between adipocytes and other cell types including stromal and endothelial cells. **d**. Comparison of cell-type deconvolution accuracy across methods using Lin’s concordance correlation coefficient (CCC), root mean squared error (RMSE), and Jensen-Shannon divergence (JSD). **e**. Comparison of estimated versus reference cell-type proportions across methods. Each point represents a cell type, with grey dashed lines indicating the identity line. **f**. Non-negative matrix factorization (NMF) of the canonical and unattributed expression component inferred by RESCUE. Weight values are min-max normalized separately for each panel. **g**. Spatial expression patterns of adipocyte marker genes, shown separately for canonical and unattributed components. Expression values are min-max normalized separately for each panel.

In adipocyte-rich regions, existing deconvolution methods fail to account for the missing adipocyte transcriptome and instead misattribute the signal to other reference cell types, such as stromal and endothelial cells (**Fig. 4b**). This systematic misassignment likely arises from transcriptional similarity, as shown by the strong correlation between adipocyte and stromal/endothelia profiles in pseudo-bulk measurements from Visium and scFFPE-seq (**Fig. 4c**). In contrast, RESCUE explicitly models unattributed expression and thus avoids forcing unassigned signal into existing reference cell types, which results in more accurate and biologically plausible estimates. As shown in **Fig. 4d-e**, RESCUE’s estimated proportions closely align with the scFFPE-seq composition, while other methods exhibit significant distortions. Together, these results demonstrate that RESCUE improves deconvolution accuracy in settings with incomplete references by preventing the systematic misassignment of unattributed transcriptomes to incorrect cell types.

To further interpret the biological content of the unattributed expression recovered by RESCUE, we applied NMF to the unattributed component. This analysis revealed multiple latent factors capturing structured expression patterns that are not explained by the canonical component (**Fig. 4f**). Notably, the NMF Factor 1 (Unattributed) captures the spatial distribution of adipocyte-rich regions, effectively recovering a missing cell type absent from the reference. Genes with the highest loadings in this factor include well-established adipocyte markers such as *FABP4, CFD*, and *G0S2*^5,56^ (**Fig. 4g**). In addition, the NMF Factor 2 (Unattributed) is enriched for genes such as *DCN, COL14A1*, and *C1R*, which have been implicated in extracellular matrix (ECM) domains^57–59^. Together, these findings indicate that the unattributed component captured by RESCUE is biologically meaningful, thereby supporting that explicitly modeling it leads to improved deconvolution in the presence of incomplete references.

Finally, to assess whether these deconvolution improvements generalize beyond the human breast cancer example, we benchmarked RESCUE using additional datasets, including human dorsolateral prefrontal cortex (DLPFC)^60^ and cichlid fish telencephalon^61^ Visium samples. In both datasets, reference profiles were derived from snRNA-seq, which inherently underrepresent cytoplasmic and process-localized RNA, leading standard deconvolution methods to misattribute unexplained ST signals. Across these datasets, RESCUE consistently outperformed competing methods (**Supplementary Fig. 3-4**), indicating that the benefits of explicitly modeling unattributed expression extend across tissues, species, and reference modalities.

### RESCUE reveals unattributed biological processes in diverse tissues

In many neuroscience experiments, cell-level information is often characterized using snRNA-seq^62,63^, which can leave a substantial fraction of biologically relevant expression unattributed. Here, we illustrate how RESCUE can uncover unique biological processes within the unattributed expression of the human DLPFC and cichlid fish telencephalon, using snRNA-seq as a reference.

#### Human DLPFC

Maynard et al.^60^ profiled the DLPFC of neurotypical adult donors using paired snRNA-seq and 10x Visium ST. We applied RESCUE to eight samples in which all six cortical layers and the white matter (WM) were annotated, to recover unattributed expression patterns not represented in the snRNA-seq reference (**Fig 5a**). As shown in **Fig. 5b**, RESCUE reveals that Layer 1 and the WM contain substantially higher levels of unattributed expression compared to other cortical layers. This is consistent with the fact that Layer 1 has sparser cell densities and contains neuropil-rich regions^57^, and that WM contains a substantial amount of axons and myelin^64^, whose transcriptomic content is mostly cytoplasmic and thus is absent from the reference snRNA-seq data.

**Figure 5.**
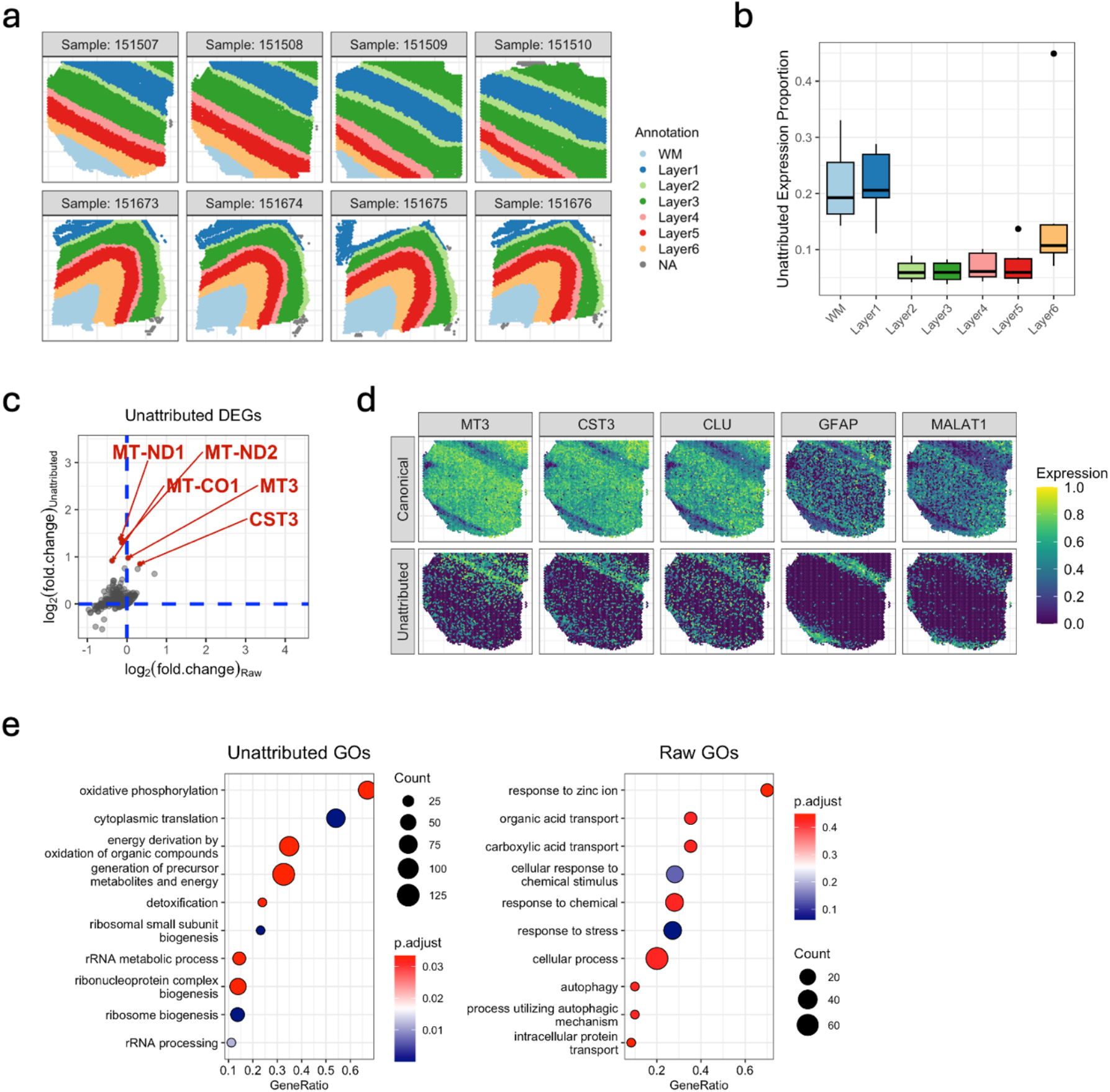
Application to the human DLPFC Visium data. **a**. Anatomical annotations of Visium ST samples from the DLPFC dataset, with cortical layers (Layer 1-6) and white matter (WM) labeled. **b**. Proportions of unattributed expression estimated by RESCUE across cortical layers and WM. Layer 1 and WM exhibit substantially higher unattributed expression compared to other layers. **c**. Comparison of log-fold changes for differentially expressed gene (DEG) analysis comparing Layer 1 to all other layers using limma on raw observed ST expression (x-axis) versus unattributed expression recovered by RESCUE (y-axis). Blue dashed lines indicate zero-fold change. Genes highlighted in red are top Layer 1 markers, detected from unattributed expression. **d**. Spatial expression patterns of Layer 1 marker genes derived from unattributed expression, shown separately for canonical and unattributed components. Expression values are min-max normalized separately for each panel. **e**. Gene ontology (GO) terms enriched among Layer 1-enriched DEGs based on unattributed (left) and raw (right) expression.

RESCUE enables us to study biology specific to the unattributed transcriptome of Layer 1. We first identified DEGs by comparing Layer 1 to all other layers using limma^65^, based on either the raw observed ST expression or the unattributed expression recovered by RESCUE. These two analyses gave substantially different results (**Fig. 5c**). For example, the top five genes most enriched in Layer 1 based on unattributed expression had either fold-changes in the opposite direction (*MT-ND1, MT-ND2*, and *MT-CO1*) or negligible fold-changes (*MT3* and *CST3*) in the differential expression analysis using raw data. We also identified genes such as *CLU, GFAP*, and *MALAT1* that exhibit clear spatial enrichment within the unattributed component of Layer 1 (**Fig. 5d**). These genes are associated with local translation and neurite outgrowth, and their enrichment outside nucleus is consistent with known roles of astrocytic processes and non-somatic cellular structures in Layer 1 of the cortex^66,67^.

To explore the underlying biological processes, we performed gene set enrichment analysis separately for DEGs derived from the unattributed expression and from the raw ST data (**Fig. 5e**). Notably, gene ontology (GO) terms enriched among the unattributed DEGs include mitochondrial processes: “oxidative phosphorylation”, “energy derivation by oxidation of organic compounds”, and “generation of precursor metabolites and energy”. In contrast, no mitochondria-related pathways reached statistical significance in the analysis based on observed ST expression; only the broad term “response to stress” passed the FDR < 0.05 threshold. These results align with the elevated metabolic demands of a neurite-enriched Layer 1, which is densely populated with synapses and reliant on local energy production to support axonal and dendritic activity^68,69^. Collectively, these findings demonstrate that RESCUE can uncover previously obscured yet functionally important gene expression programs in Layer 1 of the human DLPFC.

#### Cichlid Fish Telencephalon

Johnson et al.^61^ collected paired snRNA-seq and Visium ST data from the dorsal telencephalon of individual cichlid fishes exhibiting distinct social behaviors. As shown in **Fig. 6a**, the dorsal telencephalon is characterized by prominent radial glial (RG) processes that originate in the ventricular zone and extend long projections toward superficial pallial regions^70,71^. These RG projections span nearly the entire depth of the pallium and may play a role in shaping spatial gene expression profiles across distinct layers and compartments, but they do not survive tissue dissociation and thus their transcriptomes are not captured by snRNA-seq.

**Figure 6.**
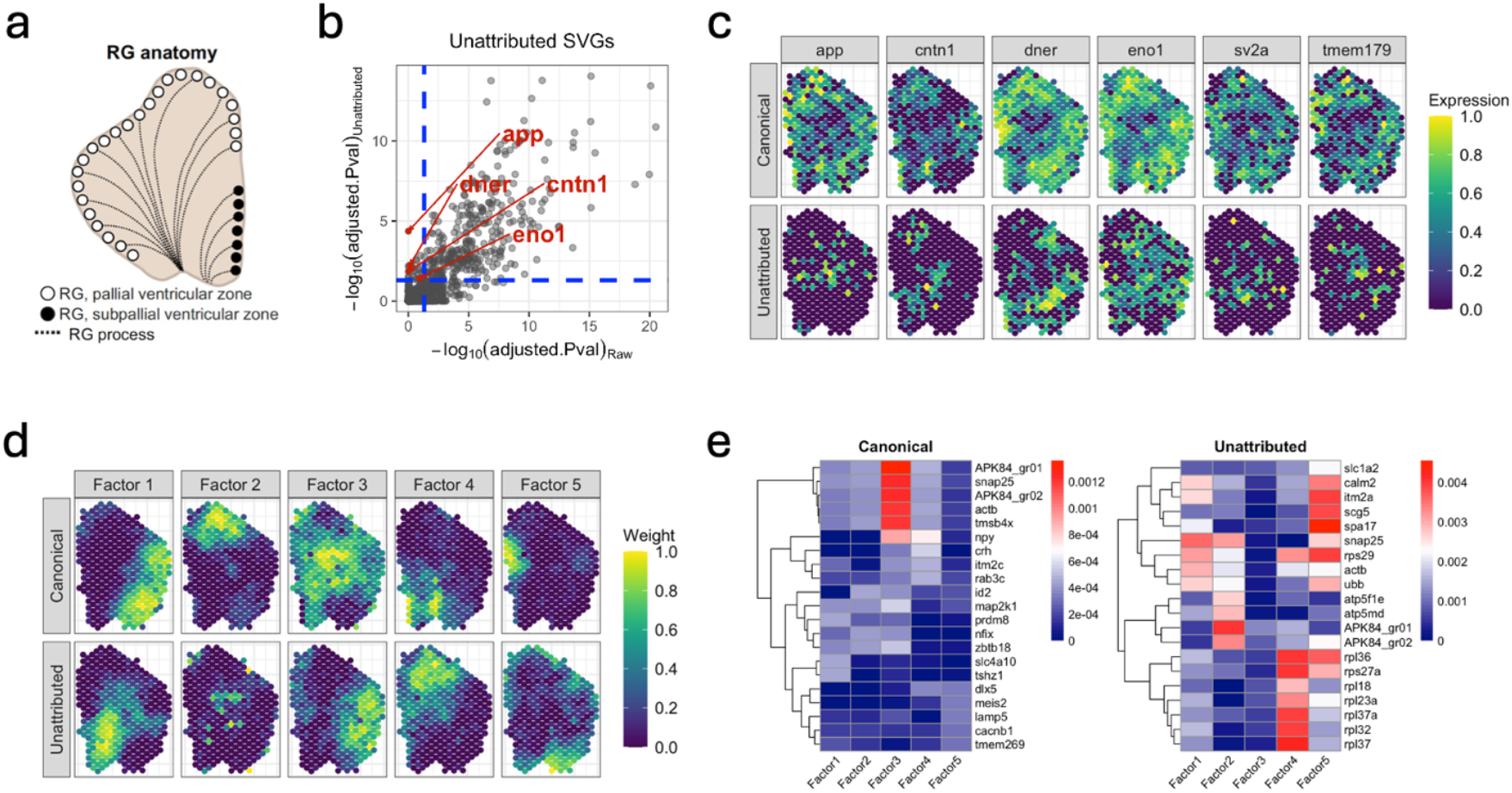
Application to the cichlid fish telencephalon Visium data. **a**. Schematic illustration of radial glial (RG) anatomy^61^ in the dorsal telencephalon, highlighting RG cell bodies in the ventricular zone and long processes extending toward superficial pallial regions. **b**. Comparison of adjusted p-values for spatially variable gene (SVG) detection using SPARK-X on raw observed ST expression (x-axis) versus unattributed expression recovered by RESCUE (y-axis). Blue dashed lines indicate the significance threshold at adjusted p-value = 0.05. Genes labeled in red are SVGs uniquely detected from the unattributed expression. Adjusted p-values are obtained using SPARK-X’s default option: Benjamini-Yekutieli procedure. **c**. Spatial expression patterns of unattributed-only SVGs, shown separately for canonical and unattributed components. Expression values are min-max normalized separately for each panel. **d**. Non-negative matrix factorization (NMF) of the canonical and unattributed expression component inferred by RESCUE. Weight values are min-max normalized separately for each panel. **e**. Heatmaps of genes with high loadings in canonical (left) and unattributed (right) factors.

We applied RESCUE to this dataset to recover unattributed transcriptomic signals using the snRNA-seq data as a reference. To assess spatial heterogeneity, we performed SVG analysis using SPARK-X^72^ on both the observed ST expression and the unattributed expression. As shown in **Fig. 6b**, a substantial number of SVGs are uniquely identified in the unattributed transcriptome. In **Fig. 6c**, we show the expression maps of those SVGs that were undetectable in the observed ST expression yet exhibited clear spatial structure once nuclear signals were removed. Among these, *app, cntn1, dner*, and *eno1* mark distinct transcriptional programs that may operate along RG projections. These genes highlight transcriptional signals that are likely to reflect localized molecular activities along RG scaffolds: *app* encodes a transmembrane protein that promotes synapse formation and plasticity^73,74^, *cntn1* (Contactin-1) is a GPI-anchored adhesion molecule essential for axon guidance and neurite outgrowth^75^, *dner* mediates neuron-glia interactions that are critical for astrocyte differentiation^76^, and *eno1* (*α*-enolase) promotes neuritogenesis and contributes to structural remodeling in the developing nervous system^77^. Indeed, Johnson et al.^61^ found that bower-building behavior in cichlid fishes may be related to neuronal rebalancing driven by a subpopulation of RG cells and their processes, highlighting the importance of studying transcriptional processes in RG projections, which RESCUE enables.

To further characterize the structure of the unattributed transcriptome, we applied NMF to the unattributed expression component (**Fig. 6d**). Several unattributed factors are enriched for genes associated with synaptic and process-localized functions, including *slc1a2, calm2*, and mitochondrial ATP synthase (**Fig. 6e**), consistent with transcriptional activity in extended RG scaffolds and neuronal processes. In addition, one factor shows strong enrichment for ribosomal protein genes, suggesting localized protein synthesis within these structures^78^. Notably, these structured transcriptional programs emerge only after the canonical expression is removed, indicating that the unattributed expression recovered by RESCUE captures biologically meaningful signals associated with radial glial and cytoplasmic processes that are not represented in snRNA-seq reference data.

## DISCUSSION

RESCUE is the first method designed to acknowledge and recover the unattributed transcriptome, which encompasses gene expression patterns not captured by cell segmentation or deconvolution approaches in ST data analysis. These unattributed signals, though typically overlooked, can carry substantial functional and biological significance. While the field has developed numerous transcriptomic profiling methods, such as scRNA-seq for isolating gene expression at the single-cell level, Ribo-seq for ribosome-associated transcripts, and CLIP-seq for binding-protein interactions, each of these technologies inevitably captures only a fraction of the total transcriptome. RESCUE provides a systematic framework to quantify and analyze what is missing: the molecular signals outside the scope of reference datasets.

By formulating the problem as a robust sparse recovery framework, RESCUE decomposes observed ST data into a reference-aligned canonical component and a biologically meaningful unattributed component. RESCUE focuses on characterizing what is not explained by reference profiles and introduces a broader conceptual innovation: computational negative selection. Rather than focusing solely on what to include (as in positive selection), we define a reference component to subtract out and focus on what remains. In this work, we used scRNA-seq and snRNA-seq data as reference profiles and focused primarily on brain tissue, but the concept is general and can be applied to a wide range of other settings, for example, muscle tissue, where multinucleated cells make segmentation challenging and single-cell reference data difficult to interpret. The choice of reference therefore requires careful design and quality control.

One striking finding emerging from our brain tissue analyses is the substantial amount of expression (26.5% of total expression) found in the unattributed transcriptome of the honey bee brain MERFISH data. While localization of mRNAs to neurites is a well-studied phenomenon^26–30^, our results, along with the estimates for the adult mouse brain from Ament and Poulpoulos^22^, additionally suggest that these local expression patterns may be different from those found in cell soma. RESCUE enables a reexamination of existing datasets to more carefully study this apparently divergent expression. Despite these promising biological insights revealed by RESCUE, reliance on computational inference alone may have potential limitations, underscoring the value of experimental validation when feasible. Orthogonal approaches, such as *in situ* hybridization, could further strengthen and refine these findings; however, such validation is often impractical in tissues where cell bodies and non-cell-body structures are spatially intermingled and difficult to segment. These challenges highlight the importance of discovery-oriented computational frameworks like RESCUE, which are designed to uncover biologically meaningful patterns and guide interpretation in settings where experimental confirmation is difficult.

There are several potential directions for future work. First, in cases where no suitable reference is available, extensions of RESCUE to reference-free or unsupervised frameworks may be possible. For example, methods based on robust principal component analysis (PCA)^79,80^ could be adapted to extract unattributed structure without predefined reference matrices. Second, applying RESCUE to high-resolution ST data requires creating a predefined grid, and further research is required to develop a systematic method to determine the optimal grid size. Third, while the gene-level sparsity assumption underlying RESCUE is biologically plausible in many settings, there may exist scenarios in which the unattributed component is not sparse but instead exhibits alternative structure, such as low-rank or low-rank plus sparse residuals. Extending RESCUE to accommodate such residual regimes represents an important direction for future research. Despite these remaining challenges, we have already shown that RESCUE can still provide meaningful analyses of unattributed expression.

Finally, RESCUE reminds us that the biology we wish to study and the measurements we are able to make are not always one and the same. For instance, we typically accept single-cell reference data as truly representing the expression of the entire single cell, but in many cases it does not, as we have illustrated above. Any measurement technology has inherent biases. As a result, recognizing, acknowledging and properly interpreting what is missed, in other words, the “unattributed expression”, is important for a comprehensive understanding of a system.

## METHODS

### RESCUE model

We assume that our ST data profiles *G* genes in *J* grids across the tissue section. In low-resolution ST technologies, these grids are predefined as multicellular spots, whereas in higher-resolution ST technologies, these can be constructed at any desired size, which allows inference at different biological scales. To introduce the statistical model behind RESCUE, we begin by reviewing existing linear deconvolution methods^12–16^. These methods decompose the ST expression matrix 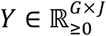 of *G* genes in *J* grids as

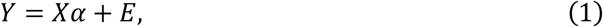

where 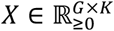 is a matrix of *K* cell-type expression reference profiles, 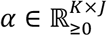 is a matrix of inferred cell-type proportions within each grid, and *E* ∈ ℝ^*G*× *J*^ is assumed to be a mean-zero noise matrix. The issue is that (1) implies that expectation of ST expression is 𝔼(*Y*) = *Xα*, which ignores unattributed transcriptome.

Instead, we further decompose the error *E*, motivated by the robust regression framework^45–47^. Specifically, we consider the multivariate mean-shift model:

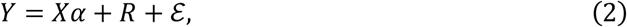

where *Xα* represents the expected expression values of each gene in each grid due to the presence of cells whose types are represented in the reference sc/snRNA-seq data, 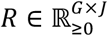 is an unattributed term that allows for outliers that cannot be captured by the reference data, and ℰ ∈ ℝ^*G* × *J*^ is a noise matrix that allows for residual variability between the ST sample and the reference data.

Following the mean-shift model in the robust regression framework, we assume that the unattributed component *R* is sparse. Specifically, the sparsity assumption reflects the expectation that only a subset of genes exhibits unattributed expression within each grid. This assumption is required for identifiability of the decomposition and is biologically plausible, as we expect relatively few genes to exhibit unattributed expression within each grid. Consistent with this assumption, we observe empirically that unattributed expression in neuropil regions of the honey bee brain is highly sparse, with 92% of genes showing no unattributed expression on average. Moreover, even in extracellular environments, ECM proteins typically involve highly sparse, restricted gene sets, with only 1.3–1.5% of genes showing nonzero expression, rather than transcriptome-wide activation^81^.

### Optimization problem

RESCUE aims to simultaneously recover the sparse non-negative matrix *R* and non-negative cell-type proportions *α* in our model (2). Specifically, we consider the following optimization problem:

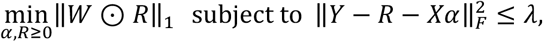

where *W* is a weight matrix, ⊙ denotes the Hadamard product, ‖·‖_*F*_ denotes the Frobenius norm, and *λ* > 0 is a tuning parameter. We set each entry of the weight matrix as 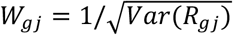 to account for the fact that different genes have different variances across different locations. These weights will be estimated through an iterative variance estimation procedure. The ℓ_1_ objective function enforces sparsity of our estimated *R*, while the constraint ensures that the residual variability ℰ is not too large, in other words, that the ST data do not deviate too much from the reference sc/snRNA-seq data.

In practice, we solve the Lagrangian form of this problem:

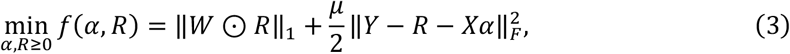

where *μ* > 0 is a tuning parameter. We apply an efficient alternating minimization to solve two convex sub-problems in each iteration:

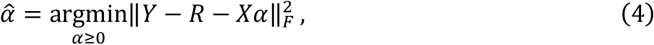

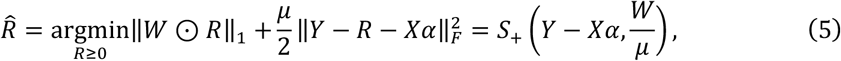

where *S*_+_ (*x, τ*) = max (*x* − *τ*, 0) denotes the entry-wise non-negative soft-thresholding function.

### Estimation of adaptive weights

Kong et al.^47^ proposed adaptive lasso-type weights to reduce masking and swamping effects in the presence of multiple outliers. Our weighting scheme plays a similar role, but differs in that the weights are learned iteratively through variance estimation of the unattributed component. In ST deconvolution models, the observed count 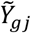 is often modeled under a Poisson assumption^13,16^:

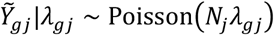

where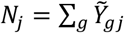. represents the total transcript count, or number of UMIs for grid *j*, and *λ*_*gi*_ denotes the expected expression rate of gene *g* at grid *j* . Rather than directly modeling 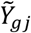, we assume that it can be decomposed as 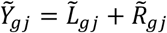, where

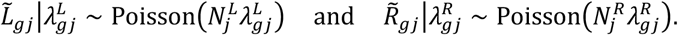

Here, 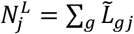 and 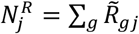 represent the total transcript counts of the canonical and unattributed expression for grid *j*, respectively, and 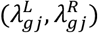 denote the expected expression rates of these components. We can write 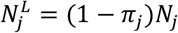 and 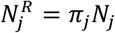, where *π*_*j*_ ∈ [0, 1] denotes the proportion of unattributed expression for grid *j*.

When using counts per million (CPM) normalization, i.e., 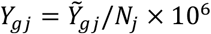, we can estimate the variance of unattributed component as

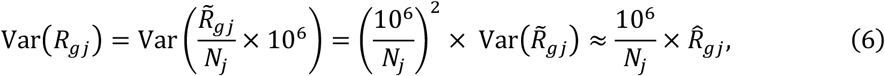

where the approximation follows from 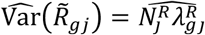 and 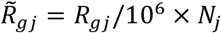. Here, the term 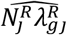 can be iteratively updated using the plug-in estimator 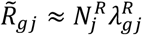. To avoid division by zero, we set 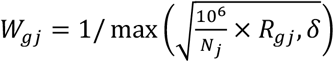, where *δ* is a small positive constant. We outline the iterative algorithm for RESCUE below (**Algorithm 1**).

#### Algorithm 1

Iterative algorithm for RESCUE

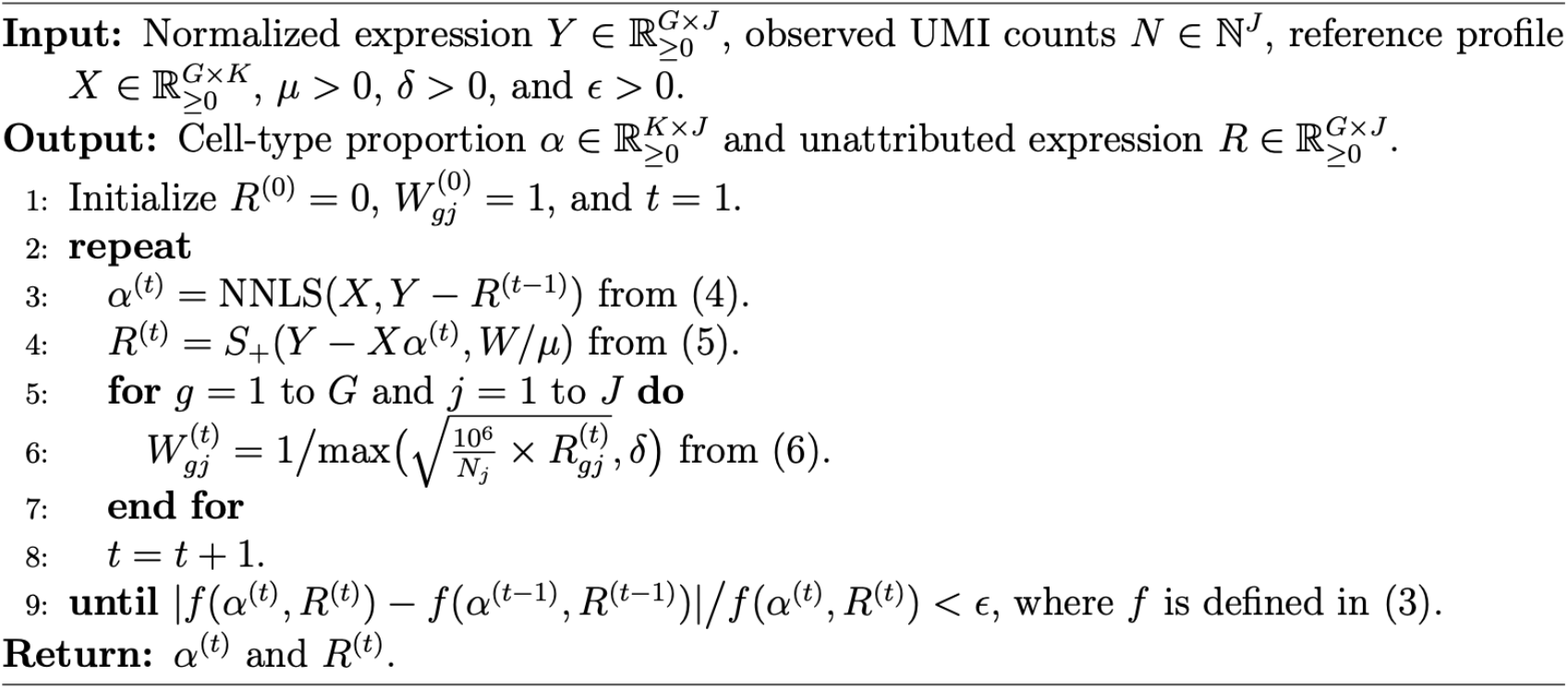

### Tuning parameter selection

RESCUE aims to recover unattributed expression by balancing the sparsity of *R*, which reflects the number of rejected outliers, against the level of residual variability ℰ. Following standard practices in robust regression and robust PCA^45,79^, we set *μ* = *GJ*/*C*‖*Y*‖_1_ and focus on selecting the optimal value of the tuning parameter *C*. However, choosing the optimal tuning parameter is challenging in the presence of outliers^80,82^.

To select the optimal tuning parameter, we adopt a data-driven strategy recently proposed for related deconvolution problems in bulk RNA-seq anaysis^83^. The key idea is to generate simulated data that closely resemble the observed ST data and use this for evaluation. Specifically, we use a surrogate ST dataset with “ground truth” proportions of canonical and unattributed expression. To construct this surrogate ST dataset, we first apply NNLS to the observed ST data to obtain initial estimates for *α* and *R*:

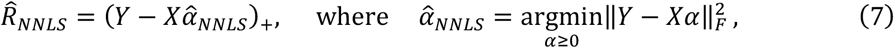

where (*x*)_+_ = max(*x*, 0) denotes the zero-thresholding function. We then simulate a surrogate ST data 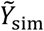 from a Poisson distribution with rescaled expected expression rates, preserving the overall signal-to-noise characteristics observed in the real data. RESCUE is applied to 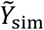 across a range of *μ* values, and the optimal *C* is selected as the value that minimizes the recovery error of the canonical and unattributed components in the surrogate dataset. We summarize the tuning procedure below (**Algorithm 2**).

#### Algorithm 2

Tuning parameter selection via surrogate match

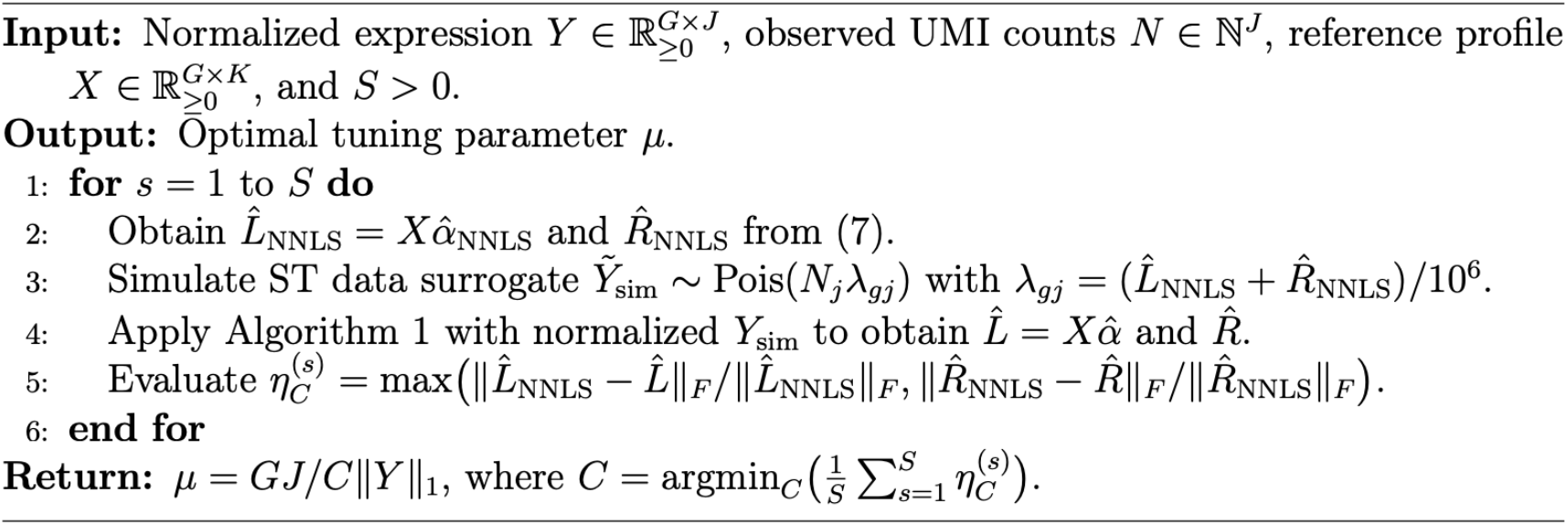

### Platform effect normalization

To mitigate cross-platform effects between ST data and sc/snRNA-seq reference data, we apply platform effect normalization. This step is essential because differences in experimental platforms can introduce systematic shifts in gene expression levels that confound downstream analysis. Without proper alignment of scale, these differences may lead to biased estimation of cell-type contributions or unattributed signals. We adopt the normalization strategy from RCTD^13^, which rescales each gene in the reference data to match the overall expression level of the corresponding gene in the ST data. Specifically, it first computes pseudo-bulk measurements for both ST and reference datasets and then apply a gene-wise log-scale correction to harmonize them.

### Data normalization and reference matrix

To account for technical variability and the effect of library size, we adopt CPM normalization but fit RESCUE in the linear space. We avoid log-transformation for two main reasons. First, log-transformed input tends to underestimate true signals in gene expression deconvolution^84^. Second, log-transformation requires adding pseudo-counts, which can introduce significant artificial distortion between zeros and non-zeros^85,86^. Instead, we work in the linear scale, employing an iteratively reweighted algorithm (**Algorithm 1**) to stabilize variance adaptively. This approach allows the model to adjust for varying data scales and minimizes the influence of outliers without artificially altering data distributions. The normalized reference matrix is then used to construct average cell-type expression.

### Reference dependence of unattributed expression

RESCUE is a computational negative selection method which relies on the choice of reference, and interpretation of the recovered unattributed expression therefore requires consideration of both biological context and reference completeness. To further assess the impact of reference selection within the same tissue, we extended the honey bee brain analysis using snRNA-seq^87^ and segmented MERFISH cells as alternative reference profiles (**Supplementary Fig. 5**). Using segmented MERFISH cells produced unattributed expression patterns highly consistent with those obtained using scRNA-seq, as both references capture cellular transcripts beyond the nucleus and therefore explain similar fractions of the ST signal. In contrast, the snRNA-seq reference yielded higher levels of unattributed expression on average, reflecting its enrichment for nuclear RNA and underrepresentation of cytoplasmic and neuritic transcripts. These results indicate that variation in the recovered unattributed component reflects biologically meaningful differences in reference coverage rather than technical artifacts, and that interpretation of unattributed expression should be guided by the biological properties of both the tissue and the reference dataset.

### Choice of grid resolution for honey bee brain MERFISH data

For high-resolution ST platforms such as MERFISH, RESCUE operates on spatially aggregated expression profiles defined over user-specified grids. The choice of grid size reflects a trade-off between spatial resolution and sparsity: smaller grids preserve fine spatial structure but increase sparsity, whereas larger grids reduce sparsity at the cost of spatial detail. As a practical guideline, we recommend selecting a grid size such that the average total transcript count per grid is comparable to the typical total transcript count per cell. In the honey bee MERFISH analysis, we used a grid size of 20 × 20 *μm*^2^, which approximates cell-scale resolution. To assess sensitivity to grid resolution, we varied grid size across a range of spatial scales and found that RESCUE produces consistent results across a broad range of grid resolutions (**Supplementary Fig. 6**).

### Subtraction-based estimation of unattributed expression for existing methods

Most existing ST deconvolution methods are designed to estimate cell-type proportions, rather than to recover expression levels directly. Because these methods do not explicitly model unattributed expression, we adopted a subtraction-based framework, following a strategy used in related deconvolution settings for bulk RNA-seq data^23^. Specifically, we first estimated cell-type proportions using each deconvolution method and then recovered the unattributed expression by subtracting the estimated cellular contributions from the observed expression, followed by zero-thresholding to avoid negative values. Under this framework, the unattributed expression is defined as the portion of the observed ST expression that cannot be explained by the reference-based cellular expression implied by each method’s estimated cell-type proportions.

### Data generating process for simulation studies

For a total of *J* = 2,500 grids, we first generated the total UMI count for grid *j* from a negative binomial distribution *N*_*j*_ ∼ NB(mean = *M*, size = 1/*σ*). We generated the raw count for each gene in each grid as 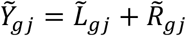, where 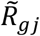 denotes the unattributed expression and 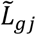 represents the canonical expression that could have been captured by sc/snRNA-seq reference data.

To generate 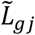, we used the real scRNA-seq data from honey bee brains^48^ as the source of canonical expression profiles. We selected the top *G* = 500 number of highly variable genes and constructed the reference matrix *X* using *K* = 20 number of cell types annotated via Louvain clustering in Seurat^88^. Latent mixing proportions were drawn as *β*_*kj*_ ∼ Beta(1,3) and normalized as *α*_*kj*_ = *β*_*kj*_/ ∑_*k*_ *β*_*kj*_, with at most 10 cell types per spot. Defining 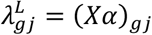, we generated the cell-type-driven canonical expression as

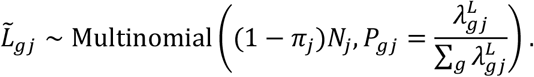

To generate sparse unattributed expression, we sampled

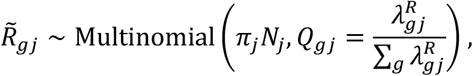

where 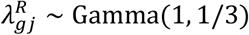 with probability (1 − *s*) and 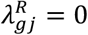 with probability *s*. Here, we introduced a sparsity parameter *s*, which makes (*s* × 100) percentage of intensity parameter 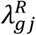 zero-inflated. Under this setting, we can generate 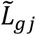 and 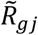 that satisfy 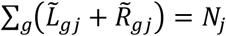. In our simulations, we fixed *M* = 10,000 and *σ* = 0.5, while varying *π*_*j*_ ∈ {0, 0.2, 0.4, 0.6, 0.8} and *s* ∈ {0.8, 0.85, 0.9, 0.95} to control the proportion of canonical expression and the sparsity level of unattributed transcriptome, respectively (**Fig. 2**). We then examined the robustness of RESCUE under additional simulation settings by varying the average total UMI count per spot (*M*), as well as varying noise levels by adjusting library size variability across spots (*σ*) (**Supplementary Fig. 1**).

### Evaluation metrics for cell-type deconvolution in simulation studies

To evaluate the accuracy of cell-type deconvolution, we considered three complementary metrics: Lin’s CCC, RMSE, and JSD. We first measured Lin’s CCC between the true and estimated proportion of the *k*th cell type, defined as

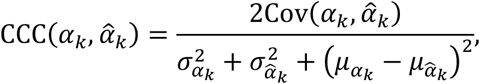

where 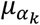 and 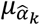 are the means of the two variables, and 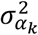 and 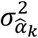 denote the corresponding variances. We used Lin’s CCC because it captures shifts in both location and scale, making it robust to high leverage points^83^. The concordance increases as the CCC value gets closer to 1. We report the average CCC across all cell types.

To quantify estimation error at the spatial location level, we computed the RMSE between the true and estimated cell-type proportion vectors. For spatial location *j*, RMSE is defined as

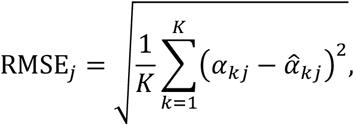

where *K* is the number of cell types, and *α*_*kj*_ and 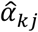 denote the true and estimated proportions of cell type *k* at location *j*, respectively. We report the average RMSE across all spatial locations, with smaller values indicating better performance.

Finally, we computed the JSD between the true and estimated cell-type proportion distributions at each spatial location. For spatial location *j*, JSD is defined as

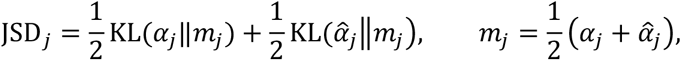

where KL(· ‖ ·) denotes the Kullback-Leibler divergence. We report the average JSD across all spatial locations, with smaller values indicating better performance.

### Evaluation metrics for unattributed expression recovery in simulation studies

To evaluate the accuracy of unattributed expression recovery, we measured the relative Frobenius error, defined as

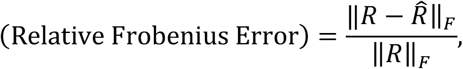

where *R* and 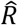 denote the true and estimated unattributed expression matrices, respectively.

### Evaluation metrics for honey bee brain MERFISH data

We measured Lin’s CCC and the RMSE between the segmentation-based (true) and estimated unattributed expression proportions, defined as follows:

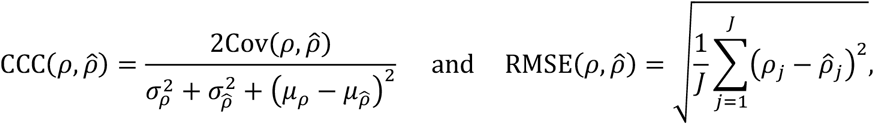

where 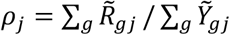 denotes the true unattributed expression proportion in grid *j*, and 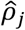 denotes its estimated value. Here, *μ*_*ρ*_ and 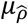 are the means of two variables, and 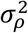 and 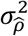 denote the corresponding variances.

### Benchmarking computational runtime

We evaluated the computational runtime of each method across ST datasets of varying sizes (**Supplementary Fig. 7**). Across all benchmarks, RESCUE completed within practical time limits for contemporary ST datasets, demonstrating scalability to datasets with hundreds to thousands of spatial locations and genes. All analyses were performed using CPU-only computation on a standard laptop (Apple M3 Pro with 36GB of memory), without GPUs or high-performance computing clusters.

## DATA AVAILABILITY

Human breast cancer dataset^5^: 10x Visium and scFFPE-seq data are available via GEO accession GSE243280. Human DLPFC dataset^60^: 10x Visium and snRNA-seq data are available at https://github.com/LieberInstitute/spatialLIBD. Cichlid fish brain dataset^61^: 10x Visium and snRNA-seq data are available via GEO accession GSE217619. Honey bee brain MERFISH data is available upon reasonable request. Honey bee brain scRNA-seq data^48^ is available at https://doi.org/10.6084/m9.figshare.16832518.

## CODE AVAILABILITY

RESCUE is available as an R package at https://github.com/brunoyjlee/RESCUE.

## ACKNOWLEDGEMENTS

We thank Dr. Tina Barbasch on valuable discussions on the analysis of fish brain data. Funding: This work was supported by the National Institutes of Health (R21HG013180 to Y.J.L., S.Y., H.-S.H. and S.D.Z., R35GM147420 to A.W.S and H.-S.H. and R01AT013189 to S.D.Z.).

